# Characterizing the Interactions of Cell Membrane-Disrupting Peptides with Lipid-Functionalized Single-Walled Carbon Nanotube Systems for Antimicrobial Screening

**DOI:** 10.1101/2023.01.25.525557

**Authors:** Anju Yadav, Payam Kelich, Nathaniel E. Kallmyer, Nigel F. Reuel, Lela Vuković

**Affiliations:** Department of Chemistry and Biochemistry, University of Texas at El Paso, El Paso, Texas, United States of America; Zymosense Inc., Ames, Iowa, United States of America; Department of Chemical and Biological Engineering, Iowa State University, Ames, Iowa, United States of America

## Abstract

Lipid-functionalized single-walled carbon nanotubes (SWNTs) have garnered significant interest for their potential use in a wide range of biomedical applications. In this work, we used molecular dynamics simulations to study the equilibrium properties of SWNTs surrounded by the phosphatidylcholine (POPC) corona phase, and their interactions with three cell membrane disruptor peptides: colistin, TAT peptide, and crotamine-derived peptide. Our results show that SWNTs favor asymmetrical positioning within the POPC corona, so that one side of the SWNT, covered by the thinnest part of the corona, comes in contact with charged and polar functional groups of POPC and water. We also observed that colistin and TAT insert deeply into POPC corona, while crotamine-derived peptide only adsorbs to the corona surface. Compared to crotamine-derived peptide, colistin and TAT also induce larger perturbations in the thinnest region of the corona, by allowing more water molecules to directly contact the SWNT surface. In separate simulations, we show that three examined peptides exhibit similar insertion and adsorption behaviors when interacting with POPC bilayers, confirming that peptide-induced perturbations to POPC in conjugates and bilayers are similar in nature and magnitude. Furthermore, we observed correlations between the peptide-induced structural perturbations and the near-infrared emission of the lipid-functionalized SWNTs, which suggest that the optical signal of the conjugates transduces the morphological changes in the lipid corona. Overall, our findings indicate that lipid-functionalized SWNTs could serve as simplified cell membrane model systems for pre-screening of new antimicrobial compounds that disrupt cell membranes.

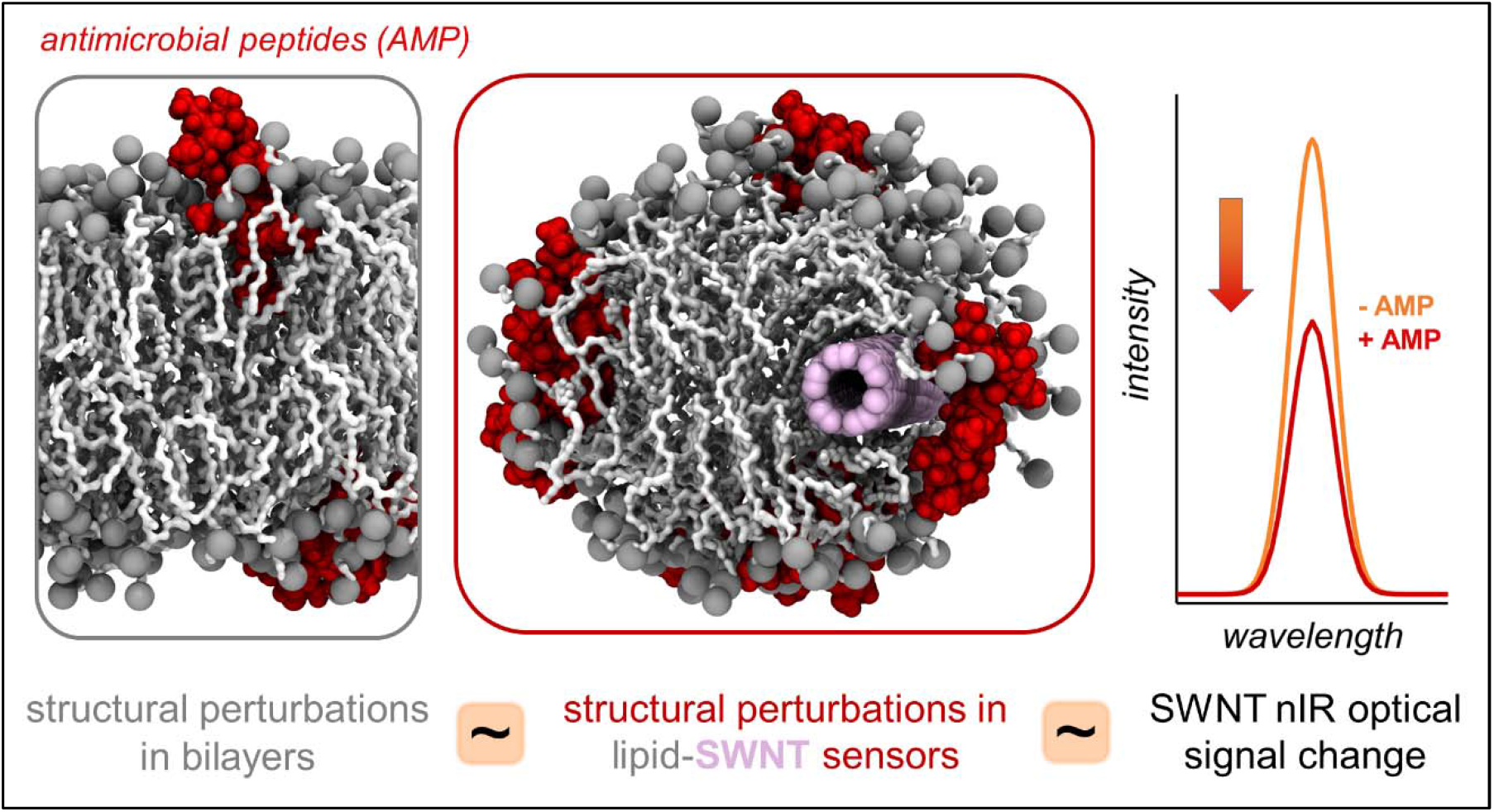

## Introduction

Many antimicrobial molecules kill bacteria by permeating and disrupting their membranes^1^. Such antimicrobials can kill bacteria cells by adsorbing to and dissolving their membranes, or by creating pores in membranes, which lead to cytoplasmic leakage and destruction of membrane potential^2,3^. Since bacteria exposed to antimicrobials naturally develop resistance over time, combatting this resistance requires continual discovery of new antimicrobials. Many organisms naturally produce antimicrobial peptides (AMP) as a part of their innate immune responses^4^. Inspired by these naturally occurring peptides, many antimicrobial screening campaigns aim to discover new membrane disrupting molecules by chemical design and synthesis of new peptides^5^ or by preparing large combinatorial libraries of peptides^6^ and testing of their antimicrobial activity. While advances in automated peptide synthesis technologies^7–9^ enable fast and inexpensive synthesis of new peptide-based compounds, testing of their antimicrobial properties still represents a major challenge.

The traditional way to screen for antimicrobial compounds is to determine their effects on live target cells using growth inhibition and cell death assays^10^. However, given the time, financial costs, low throughput, and the contamination risks associated with using live cell assays, other helpful antimicrobial screening approaches have been developed for antimicrobial discovery and optimization. Droplet-based microfluidic platforms have been developed, where single cells and candidate compounds can be encapsulated into picolitre-volume emulsion droplets and the resulting effects analyzed by imaging, specially incorporated fluorescence reporters, or other means^11,12^. Other approaches based on abiotic systems have also been developed, which can screen for molecules that specifically disrupt cell membranes. These systems can be based on simplified cell membrane models, such as the artificial planar lipid bilayers and the liposomes^13,14^. Planar lipid bilayers can be stabilized on solid support, and the effects that candidate compounds have on these bilayers can be examined by surface plasmon resonance spectroscopy^15^. While this technique can provide important biophysical details, such as the apparent rate and affinity constants, which are useful to study the mechanisms of pore formation in membranes, it lacks the throughput required for larger screening applications. Liposomes or lipid vesicles can be prepared to contain dyes in their interiors, which can be tracked optically in time to report if candidate compounds lead to disruption of vesicle membranes and the consequent leakage of dye from the vesicle.

The existing methods for antimicrobials discovery can be helpful for certain stages of screening processes. However, each method has its own inherent advantages and limitations, the latter often related to speed and throughput, the level of detail they provide into the mechanisms of antimicrobial activity, and the applicability of results to live cells. New technologies that address some of these limitations would be helpful and could become a part of the toolbox for antimicrobial discovery efforts. Recently, an abiotic system based on lipid-wrapped single-walled carbon nanotube (SWNT) conjugates has been proposed and investigated as a new optical sensing platform for screening of antimicrobial compounds^16^. This platform uses SWNTs as near infrared (nIR) fluorescent transducers to report lipid interactions with antimicrobial compounds that take place on the SWNT surface. The authors in Ref.^16^ investigated SWNTs wrapped by 1-palmitoyl-2-oleoyl-sn-glycero-3-phosphocholine (POPC) molecules and several bacterial lipopolysaccharides and proposed that the observed optical responses to known cell-penetrating AMPs (decrease in fluorescence signal intensity) are associated with morphological changes to lipids on the SWNT surface, induced by peptide binding and lipid disruption.

SWNTs wrapped by polymers, including surfactants, lipids, single stranded DNAs and RNAs, peptides, and other molecules^17–27^ have been prepared for a variety of applications, such as biological catalysis^28^, bioseparations^29–31^, gene therapy^32,33^ photo-thermal cancer therapy^34,35^, and detection of bioanalytes^36–40^. The role of polymers, which form the corona phase on the SWNT surface, is to both solubilize hydrophobic SWNTs in the aqueous media and to bind to potential analytes of interest in sensing applications. On the other hand, SWNTs, with their exceptional electronic and optical properties^41^, are the key optical sensing components in polymer-wrapped SWNT conjugates. SWNTs fluoresce in the nIR region in the 820 nm to 1600 nm wavelength window^42^ and this fluorescence signal is photostable and does not bleach^41^. There are multiple mechanisms by which a change in the SWNT environment can alter the fluorescence emission of SWNTs. Changes in the molecules surrounding SWNTs can induce a solvatochromic effect^41,43–45^, where, for example, the change from the low dielectric constant hydrophobic chains of sodium dodecyl sulfate to the higher dielectric constant N-Methyl-2-pyrrolidone solvent around SWNTs leads to red shifts in the wavelength of SWNT fluorescence^43^. The SWNT emission can be altered by changing the local electric fields at the SWNT surface^46^, which may occur when molecules with charged functional groups bind to SWNT conjugates. Furthermore, binding of charged redox-active molecules to SWNT conjugates can also alter SWNT emission^25^. Changes in the intensity of the SWNT emission were also reported, induced by the changes in the dielectric constant of the media surrounding the SWNT, ε_r_. For example, a change in the dielectric constant around the SWNT from ε_r_ = 2 to 5 was shown to lead to the fluorescence emission intensity drop of more than 50%^45^.

While the above previous studies correlated the wavelength and intensity changes in SWNT emission to the major changes in dielectric constants and local electric fields in the SWNT environment, it is still mostly unclear how small concentrations of analytes change the SWNT environment. In case of the POPC-wrapped SWNTs binding to antimicrobial peptides, the addition of small concentrations of these peptides was shown to primarily decrease the intensity of the SWNT emission^16^. While the optical signals of lipid-wrapped SWNTs in response to addition of antimicrobial peptides were determined experimentally^16^, no structural studies of these systems have been reported to date, and therefore, the actual structural changes in lipid corona that induce the decrease in the SWNT fluorescence emission intensity are unknown. Furthermore, it is unclear if the peptide-induced structural changes in lipid-SWNTs are qualitatively and quantitatively related to the structural changes these peptides make in bilayer membranes made of the same lipids. Importantly, if the structural changes induced by antimicrobial peptides in lipid-SWNTs and lipid bilayers are similar in nature, and the optical response of SWNTs truly transduces the nature and magnitude of lipid phase structural changes, then lipid-SWNTs can be used as simplified cell membrane models for fast screening of new molecules that disrupt cell membranes.

In this study, we characterize phospholipid-coated SWNTs, investigate the effects of membrane disrupting peptides on phospholipid-coated SWNT sensors and phospholipid bilayers, and compare the observed effects in two phospholipid systems. We use atomistic molecular dynamics (MD) simulations to characterize peptide binding and perturbations of these phospholipid systems. The results of our simulations and the results of the previously performed experiments^16^ are then examined for possible correlations and mechanistic interpretations.

## Results

### Asymmetric Distribution of POPC Corona Around SWNTs

Our initial MD simulations characterized POPC lipids interacting with (6,5) SWNTs. Four systems were built with POPC lipids placed in cylindrical arrangement around SWNTs (**Figure S1, Table S1**). These four systems contained 9:1, 15:1, 20:1 and 30:1 POPC to SWNT mass density ratios, where 30:1 ratio corresponds to the mass density ratio used in previous experimental preparations^16^. Modeled systems contained single SWNTs because POPC-SWNT conjugates are presumed to emit light and thus perform sensing only when containing single SWNTs, whereas conjugates with bundled SWNTs have quenched emission^47–49^.

After 175 ns to 1 μs of equilibration in MD simulations, four prepared systems relaxed to structures shown in **Figure 1a**. While the POPC-SWNT conjugate with 9:1 mass density ratio relaxed into a cylindrical shape, other conjugates assumed bilayer-like shapes in (x-y) dimensions, with the bilayer resemblance increasing with the mass density ratio. In all four systems, SWNT centrality was lost, resulting in POPC coronas around SWNTs developing marked asymmetry over time. This asymmetry development over time for 15:1 POPC-SWNT system is shown in **Figure S2**. The equilibrated SWNTs become wrapped by hydrophobic POPC tails. However, one side of SWNT is always in contact with the zwitterionic head groups of POPC molecules. The asymmetry is evident in the measured thickness of POPC corona for systems with 9:1 and 15:1 mass density ratios, shown in **Figure 1b**. The thickest parts of the POPC corona span 3.5 nm and 3.9 nm for 9:1 and 15:1 systems, while the thinnest parts of the POPC corona are ∼0.5 nm to ∼0.6 nm wide for these systems.

**Figure 1.**
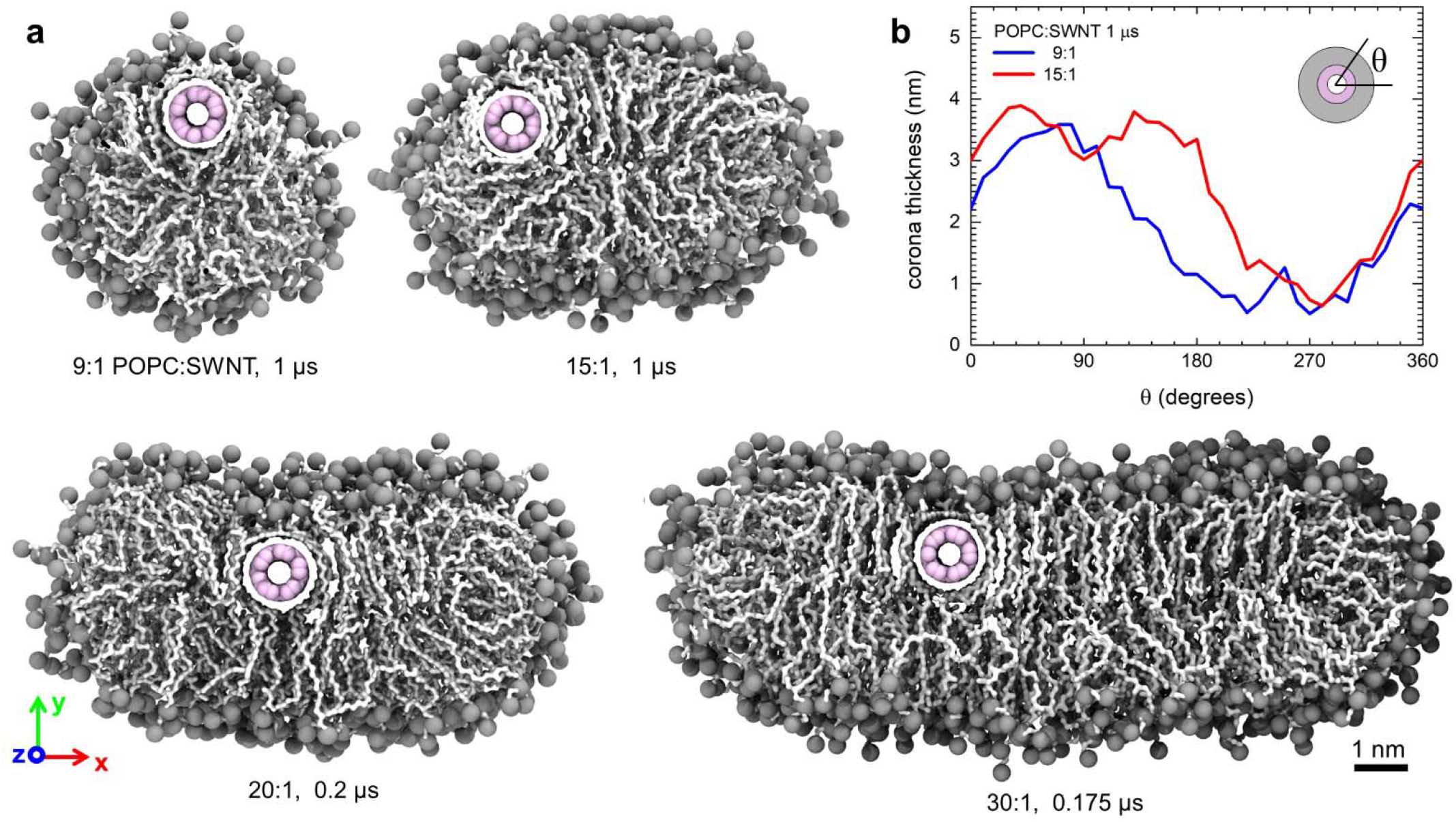
Models of POPC-SWNT conjugates equilibrated in MD simulations. a) Snapshots of four POPC-SWNT systems with 9:1, 15:1, 20:1, and 30:1 POPC:SWNT mass density ratios after the equilibration in MD simulations. Equilibration times are stated underneath the images. SWNT atoms are shown in pink, POPC tails are shown as white chains, and P and N atoms of POPC are shown as grey spheres. Water is not shown for clarity. b) POPC corona thickness around SWNTs for systems with 9:1 and 15:1 POPC:SWNT mass density ratios, measured after 1 μs of equilibration.

### Interactions of POPC Membrane Disruptors with POPC-SWNTs

Next, we examine the interactions of POPC-SWNT systems with 9:1 and 15:1 mass density ratios and three peptides, known to disrupt or bind to POPC membranes, vesicles, or POPC-SWNTs. These three peptides include colistin^50^ (Col), TAT peptide^51^ (TAT) and crotamine-derived peptide^16,52^ (Cro). After adding ten molecules to equilibrated POPC-SWNT systems, the resulting combined systems were then equilibrated in aqueous solutions for 1 μs each, resulting in the structures shown in **Figure 2a**. The disruptor molecules bind to all parts of the POPC corona, with some molecules binding to thick sections of the POPC corona, and other molecules coming in direct contact with the nanotube and thin sections of the POPC corona. Two types of binding modes are observed for molecules interacting with thick sections of the corona. In the first binding mode, some molecules insert deeply into it, especially Col and, to a lesser extent, TAT. When inserting into the corona, these molecules form many contacts to POPC, both with the hydrophobic chains and the zwitterionic headgroups of POPC. In the second binding mode, the molecules interact primarily with the zwitterionic headgroups. This second binding mode is predominantly observed for Cro, and some of Col and TAT molecules.

**Figure 2.**
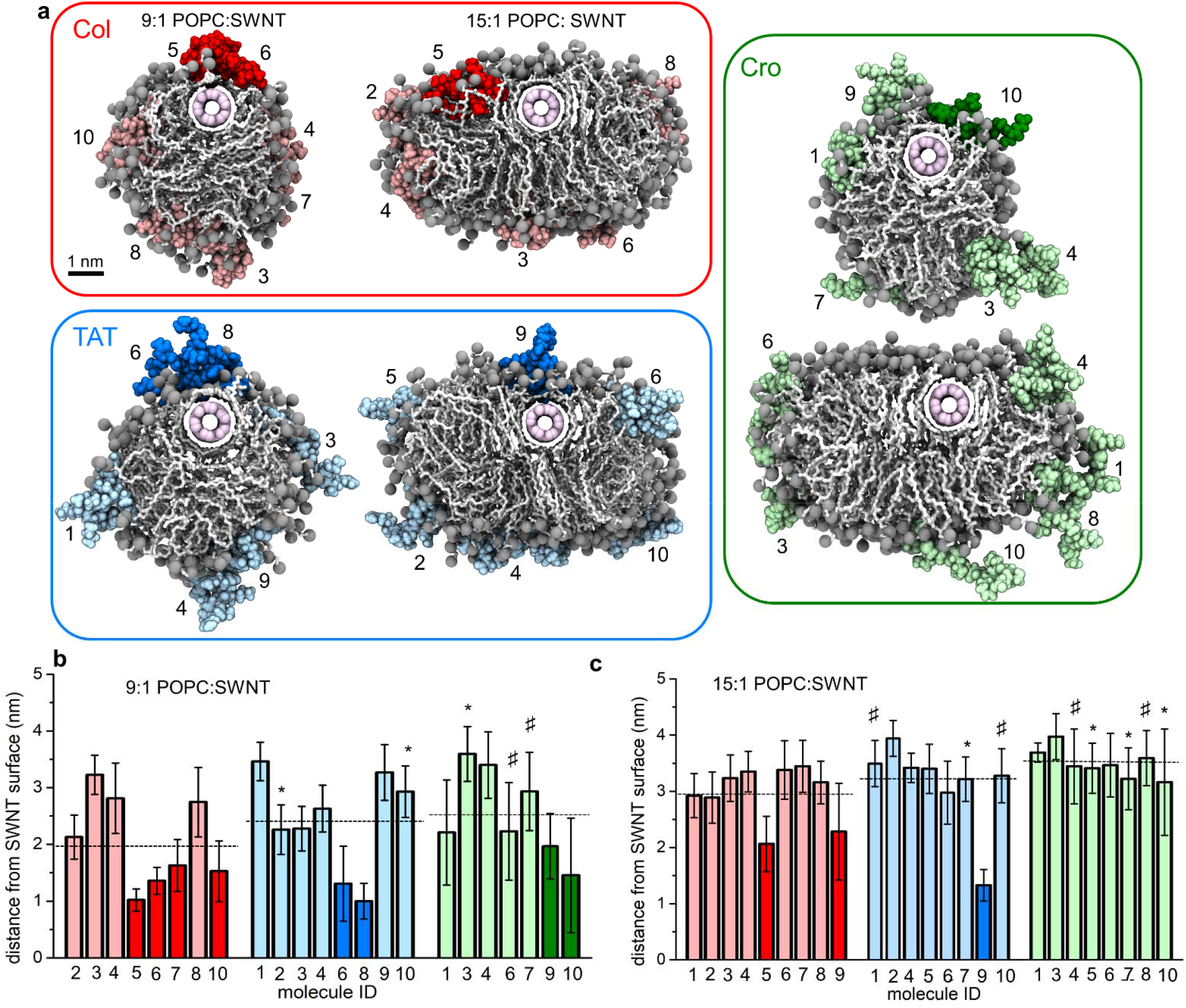
Binding of colistin, TAT peptide and crotamine-derived peptide to POPC-SWNT conjugates. a) Snapshots of equilibrated POPC-SWNT conjugates interacting with Col (red), TAT (blue), and Cro (green). The conjugates have 9:1 and 15:1 POPC:SWNT mass density ratios. Deeper hues mark molecules with direct SWNT contact in the last halves of trajectories, and lighter hues mark molecules bound to thicker parts of POPC coronas distant from the SWNT surface. b) Distances of molecules (geometric centers) from SWNT surfaces, averaged over the times the molecules are bound to POPC-SWNTs, summarized in Table S4. Histogram bar color hues distinguish the SWNT-bound (deeper hue) and corona-bound molecules (lighter hue). Bars labeled with (*) symbol indicate molecules that bind and unbind from POPC-SWNT, and bars labeled with (D) symbol indicate molecules that undergo two binding events to POPC-SWNT. Dashed lines mark average distances of all bound molecules from the SWNT surface for each system.

The plots in **Figure 2b-c** quantify the average distances of bound molecules (geometric centers) from SWNT surfaces. For both 9:1 and 15:1 systems, Col has the largest number of molecules stably bound, which never unbind from POPC-SWNT after the initial binding event. TAT molecules have the next largest number of molecules bound to POPC-SWNT, followed by Cro. However, both TAT and Cro are occasionally observed to unbind from POPC-SWNT over the course of the trajectory, as marked in **Figure 2b-c** and **Table S4**. Col and TAT each have either two molecules (9:1 systems) or one molecule (15:1 systems) binding directly to the SWNT surface, while Cro only has a single molecule binding to the SWNT surface in 9:1 system. Furthermore, in both 9:1 and 15:1 systems, bound Col molecules are on average closer to the SWNT surface, followed by TAT and then Cro. Separately, we note that disruptor molecules do not appear to influence the asymmetric distribution of the POPC corona around the SWNT. **Figure S3** shows how the thinnest part of the POPC corona around SWNT evolves over time in 9:1 and 15:1 POPC-SWNT systems with and without the added disruptors. In both types of POPC-SWNT systems, the thinnest parts of the POPC corona are of similar width, showing that disruptor molecules do not further thin out the POPC corona.

Overall, the results of **Figure 2** allow us to rank three peptides according to the magnitude of their disrupting effect on POPC-SWNT systems. Col molecules bind most stably to POPC-SWNT without unbinding, and they insert most deeply into the POPC corona on average (dashed lines in **Figure 2b-c**). Col and TAT also have the largest fraction of molecules binding directly to the SWNT surface, thus being the most likely molecules to perturb the nIR emission of SWNTs. The disruptors with the intermediate and the smallest number of bound molecules are TAT and Cro, respectively. Therefore, of the three molecules examined, Col shows the greatest disruption effect on POPC-SWNT, and the total ranking according to the disruption effect on POPC-SWNT in a decreasing order is Col, TAT, and then Cro.

### Interactions of Peptides with POPC Bilayers

Phospholipid-functionalized SWNTs could be a novel nanotechnology tool capable of discovering new molecules that disrupt membranes^16^. To have this capability, the optical signal emitted by SWNTs in the presence of candidate molecules should be proportional to the structural disruption of the lipid corona on SWNT and, importantly, to the disruption that the candidate molecules cause in the actual membrane bilayers.

Therefore, we next examined the interactions of Col, TAT and Cro with pure POPC bilayers, while holding number concentration ratios between three molecules, POPC and water the same as in the POPC-SWNT systems (Methods, **Table S2**). **Figure 3a** shows the structures of peptides binding to POPC bilayers following 1 μs equlibration in MD simulations. Compared to POPC-SWNT systems, the fraction of molecules bound to pure bilayers is lower, as only four Col, and two TAT and Cro molecules bind to bilayers at the end of the simulations. Furthermore, unbinding events are observed in bilayer systems over the course of 1 μs simulations (**Table S5**).

**Figure 3.**
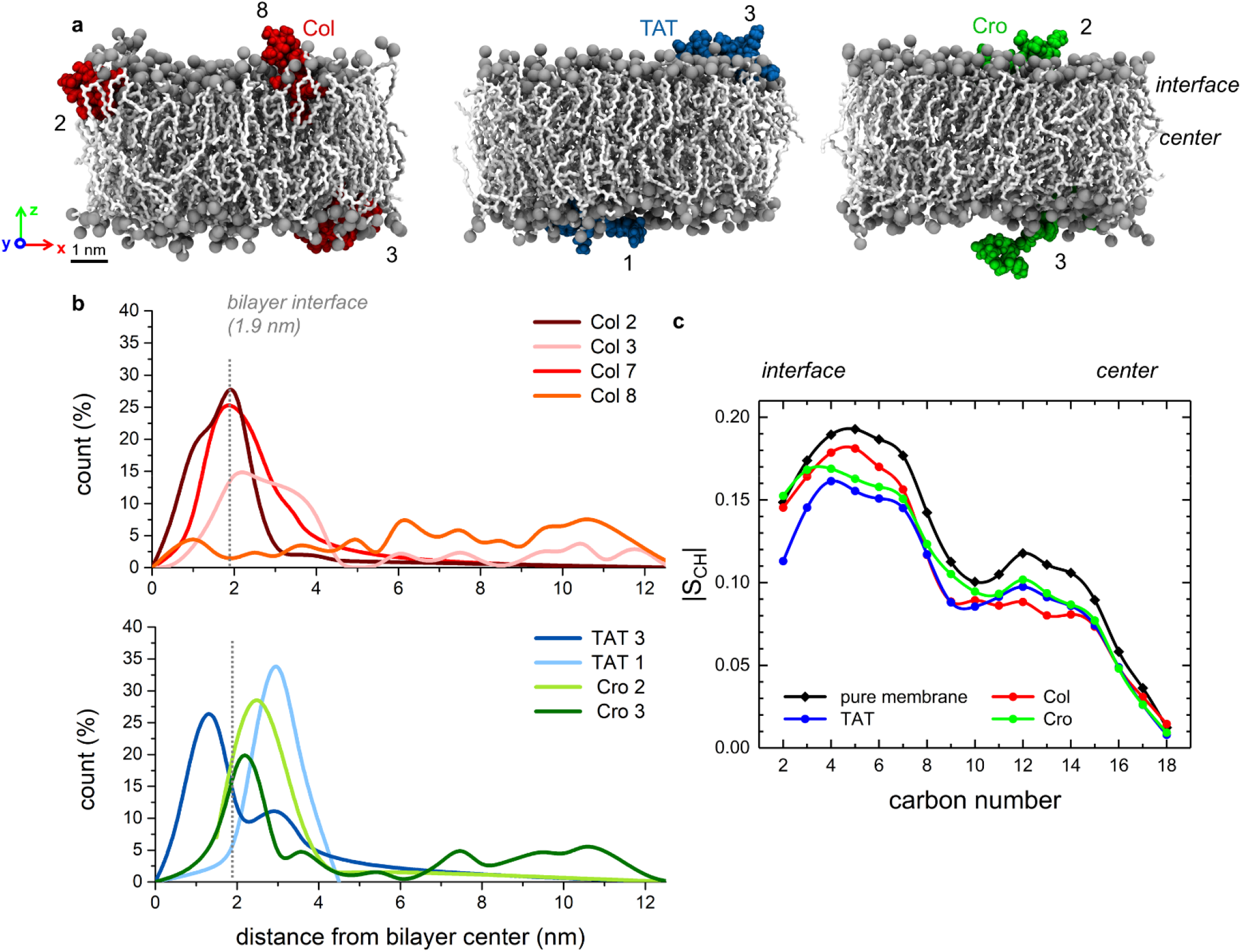
Binding of colistin, TAT peptide and crotamine-derived peptide to pure POPC bilayers. a) Snapshots of equilibrated POPC bilayers interacting with Col (red), TAT (blue), and Cro (green). The colors and representation schemes are the same as in previous figures. b) Distribution of geometric center distances (z-axis components) for bound Col, TAT and Cro molecules with respect to the bilayer center. c) Order parameter of POPC bilayers (pure and with bound Col, TAT, or Cro molecules).

Despite the binding concentration differences, there are nonetheless marked similarities between binding modes of the three molecules to POPC bilayers and to POPC-SWNTs. First, more Col molecules than TAT and Cro molecules bind to the bilayer, in agreement with the trend observed for binding of these molecules to POPC-SWNTs. Second, bound Col molecules tend to insert deeply into the bilayer (**Figure 3a**), as was also observed for Col in POPC-SWNT systems. **Figure 3b** quantifies the probability and depth of insertion into bilayers for each of the three types of molecules examined. Distribution plots for bound molecules in **Figure 3b** reveal that three out of four bound Col molecules spend a significant fraction of time inserted deeply into the hydrophobic part of the bilayer, within 1.9 nm from the bilayer center. In contrast, only one of two TAT molecules inserts deeply into the bilayer, and Cro inserts the least into the hydrophobic part of the bilayer.

Next, we tracked the magnitude of the disruption for examined molecules by calculating the order parameters of POPC chains in the simulated bilayers. First, we confirmed that our pure POPC bilayer has an order parameter profile that matches the experimentally determined order parameter profile (**Figure S4**). In general, the value of S_CH_ = -0.5 is obtained if all the lipid chains are perfectly aligned (parallel) with the membrane normal z-axis, whereas the value of S_CH_ = 0 indicates an isotropic system with completely unordered lipid chains. **Figure S4** shows that in pure POPC bilayer the atoms at the zwitterionic interface are more ordered and atoms towards the bilayer center are more disordered. Next, we examined the effect that the disruptor peptides have on the POPC bilayer order parameter. As shown in **Figure 3c**, all three peptides reduce the ordering of POPC chains, both at the zwitterionic interface and further into the bilayer core. Col is also found to disrupt the parts of the bilayer that are closer to the bilayer center.

The results shown in **Figure 3**, namely, the number of disruptor molecules bound to the POPC bilayer and their insertion depths, allow us again to rank these molecules according to their bilayer-disruption activity. The ranking of the molecules is Col, TAT, and Cro in the order of the decreasing bilayer disruption activity. This ranking matches the ranking observed for these molecules interacting with POPC-SWNTs, indicating that POPC-SWNT conjugates experience the disruption by the added molecules which is similar in nature and the extent of disruption that these molecules produce in the analogous lipid bilayers.

### Local Disruptions at the SWNT Surface and Their Correlation to the Optical Signal Modulation by Disruptor Peptides

SWNT optical emission was shown to depend on dielectric properties of the environment surrounding the SWNT^43,45,53^. In the systems examined in the present study, the environment is highly heterogeneous, including hydrophobic and ionic groups of POPC, polar water molecules, free counterions, and cationic disruptor peptides. When peptides are binding to POPC-SWNT, they can displace POPC, and modify the concentrations of water and ionic groups near the SWNT surface. Therefore, peptide binding can induce changes to the SWNT environment, and the optical emission of SWNT could be different before and after the addition of peptides. Here, in separate simulations, we next examine how the SWNT environment in the region of the thinnest POPC corona changes in response to the addition of three membrane disruptors, Col, TAT and Cro.

First, we examine the amounts of water and ions that come in direct contact with the SWNT surface and compare these amounts in pure POPC-SWNT systems and the POPC-SWNT systems in which the disruptors are bound to the thinnest part of the POPC corona. The following analyses were performed for separately simulated POPC-SWNT systems which contain only single disruptor molecules binding to the thin region of the corona, summarized in **Table S3. Figure 4a** shows a representative snapshot of one of the systems, including a TAT peptide binding to POPC-SWNT with 15:1 POPC:SWNT mass density ratio in the region where POPC corona is at its thinnest (∼0.6 nm). While most of the SWNT surface is covered by POPC chains, there is also a TAT molecule and a significant number of water molecules in direct contact with SWNT.

**Figure 4.**
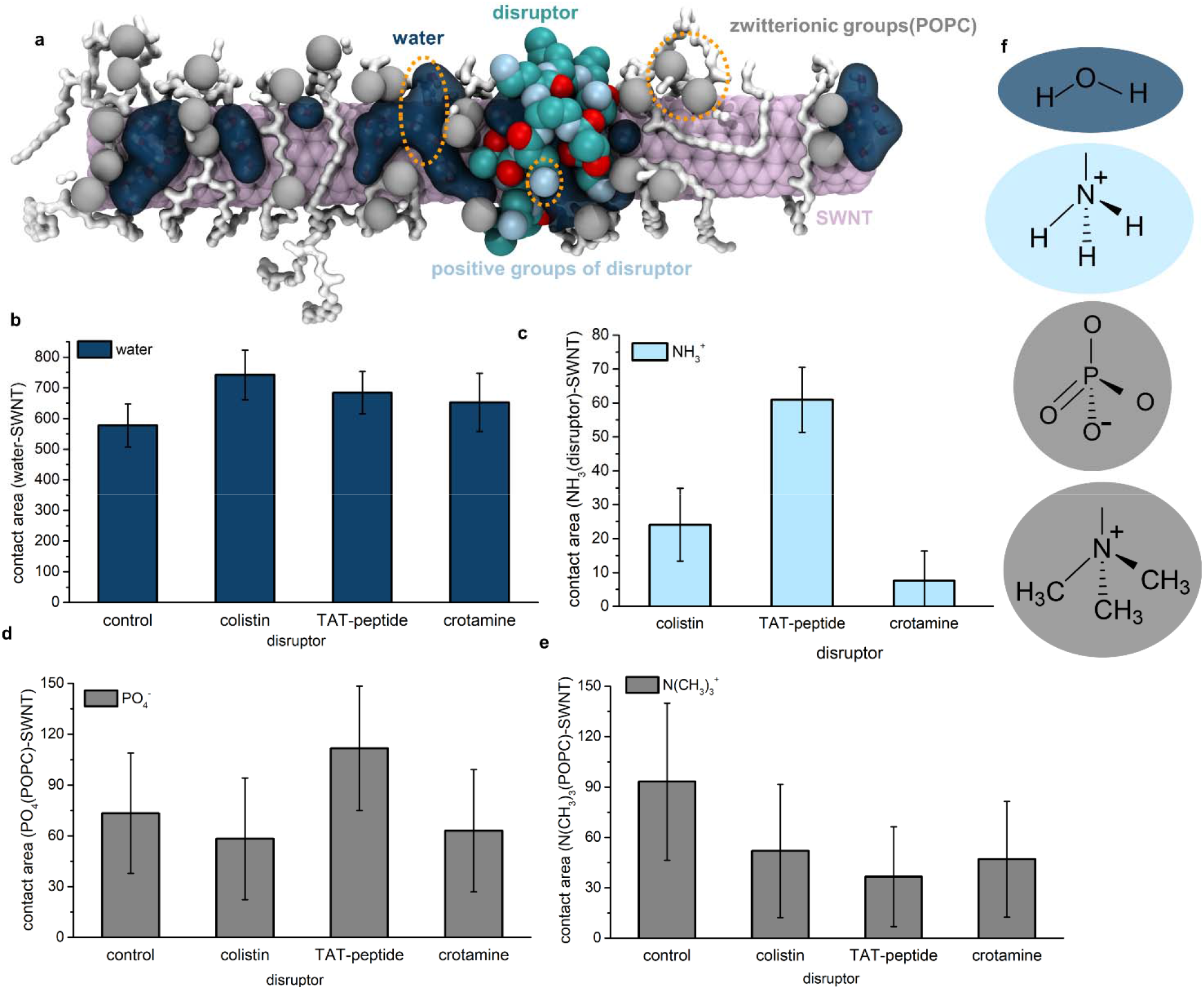
Analysis of functional groups in contact with the SWNT surface for control POPC-SWNT systems and systems interacting with Col, TAT, and Cro. a) A snapshot of a POPC-SWNT system (15:1) binding to one TAT molecule. SWNT and POPC atoms are shown as in previous figures. Water is shown as a dark blue surface and TAT atoms are shown as spheres (C – teal, O – red, and N – light blue). For clarity, the snapshot includes only the water molecules and POPC atoms near to the SWNT surface. The orange dotted ovals highlight the polar/charged functional groups, whose chemical structures are shown in panel f. b) Contact areas between water molecules and the SWNT surface. c) Contact areas between -NH_3_ ^+^ groups of disruptor molecules and the SWNT surface. d) Contact areas between -PO_4_^-^ groups of POPC molecules and the SWNT surface. e) Contact areas between -N(CH_3_)_3_ ^+^ groups of POPC molecules and the SWNT surface. f) Chemical structures of the functional groups examined in panels (b-e). The contact areas in panels (b-e) are averaged over the last half of production run trajectories.

To quantify the number of polar or charged moieties that are in direct contact with SWNT, we calculated the contact areas between SWNT and several moieties of interest, including water and charged functional groups on POPC and disruptor molecules. **Figure 4b** shows the average contact areas between water and SWNT surface in four simulated systems. In the pure POPC-SWNT system, water contacts 577 Å^2^ of the SWNT surface. The presence of disruptors leads to more water coming in contact with the SWNT surface. Namely, binding of Col, TAT and Cro molecules results in water making contact with 742, 684, and 652 Å^2^ of the SWNT surface, respectively, a 10-20 % increase in water-SWNT contact compared to the contact observed in the pure POPC-SWNT system. The trends observed in water-SWNT contact area analyses agree with the analyses of the number of water molecules in contact with SWNTs, shown in **Figure S5a**. Therefore, the addition of disruptor molecules and their binding to POPC-SWNT conjugate tends to increase the amount of polar water in contact with the SWNT surface.

All the disruptor molecules are positively charged, with Col and Cro having a +5 charge and TAT-peptide having a +8 charge in neutral pH solutions. Most of the positive charges are due to -NH_3_^+^ groups, in addition to the guanidinium groups, which contribute to the positive charge of TAT. **Figure 4c** shows the average SWNT surface area that comes in contact with NH_3_^+^ groups of the disruptor molecules. The disruptor with the largest net positive charge, TAT, leads to the highest SWNT area being in contact with the positive charge, followed by Col and Cro. Plots in **Figures 4d-e** show the average SWNT surface areas that come in contact with the charged groups of POPC, namely the -N(CH_3_)_3_^+^ and -PO_4_^-^ groups. The presence of all three peptides reduces by ∼40% the number of -N(CH_3_)_3_^+^ groups in contact with SWNT. The presence of the disruptors also leads to a similar number of -PO_4_^-^ groups in contact with the SWNT for pure POPC-SWNT systems and the systems with the disruptors, except in the case of TAT, which increases the number of the PO_4_^-^ groups near SWNT surface. Again, the trends observed in the contact area analyses agree with the analyses of the number of the charged groups in contact with SWNTs, shown in **Figure S5**. The observed trends are likely related to the magnitude of positive charge which the disruptor molecules possess, so that the positively charged disruptors repel the positive functional groups and sometimes attract more of the negative functional groups on POPC molecules to the SWNT surface.

Overall, the results of **Figure 4** and **Figure S5** show that when the disruptors are introduced to the POPC-SWNT systems, the amount of water and the number of the charged groups surrounding the SWNT surface increase. Due to the presence of a different number of polar and charged groups around the SWNT in the absence and presence of the disruptor molecules, the dielectric constant of the SWNT environment is also likely to be different in the absence and presence of these molecules. These dielectric changes upon the addition of disruptors to POPC-SWNT systems could be the primary origins of the quenching of the nIR signal, which was observed in the experiments^16^ and which was shown to be disruptor-dependent.

Next, we examined how two of the properties calculated in simulations correlate with the experimental nIR emission change of lipid-wrapped SWNTs after addition of disruptor molecules^16^. **Figure S6** shows the correlations between the experimentally measured SWNT nIR emission change and two properties from simulations, namely, the fractions of time that disruptor molecules bind to POPC-SWNT, and the water-SWNT contact area change due to addition of disruptors. In **Figure S6**, the nIR SWNT emission changes due to addition of TAT and Cro were experimentally measured for POCP-SWNT conjugates, whereas the nIR SWNT emission change due to addition of Col was measured for SWNTs wrapped by lipopolysaccharide from Escherichia coli (serotype O26, Sigma L8274)^1^. The plots in **Figure S6** show correlation between the computed properties and the experimentally measured POPC-SWNT nIR emission changes. Overall, the results show that small molecules with stronger interactions with POPC-SWNT conjugates, which also bring more water to the SWNT surface upon binding, induce a larger reduction of the SWNT nIR emission. It seems self-evident that the molecules with stronger interactions with POPC-SWNTs and which bring more water to the SWNT surface are also the molecules which induce larger disruptions in the POPC corona. As we confirmed in **Figure 3**, such molecules are also more likely to induce larger disruptions of the POPC bilayer membranes.

## Conclusion

In this work, we first characterized the POPC-coated (6,5) SWNT conjugates using MD simulations. We initially expected the equilibrated SWNTs to be solvated in centers of POPC cylindrical micelle-like coronas, between the ends of hydrophobic tails of POPC lipids. Such central positioning was observed in previous simulations of (18,0) SWNTs solubilized in other lipids and surfactants, including two detergents lysophosphatidylcholine and dihexanoylphosphatidylcholine, and a bilayer-forming lipid dipalmitoylphosphatidylcholine (DPPC)^54^. These previous 1.6 μs long simulations used coarse-grained parametrization of systems with MARTINI force field^55^, widely used to simulate lipid assemblies and their interactions with nanoscale systems^56–60^. Notably, the central positioning of SWNTs was observed for several lipid:SWNT concentration ratios and using (18,0) SWNTs^54^, which are almost twice wider than the (6,5) SWNTs used in the present study.

Contrary to our expectations, SWNTs in all our simulations became asymmetrically positioned within POPC coronas, as shown in **Figure 1**. This asymmetric non-central positioning developed during the first 50 ns of our atomistic simulations, as the SWNTs seemed to be extruded from the most disordered central region of the POPC micelle, formed by the ends of hydrophobic tails of POPC lipids. Due to the asymmetric positioning, the part of the SWNT where the POPC corona is the thinnest comes in contact with charged and polar functional groups of POPC and water. The rest of the SWNT is in contact with the hydrophobic sn-1 and sn-2 chains, some of which become parted and wrap the SWNT. Our observations resemble the results of Ref.^61^, where SWNTs solvated in lysophosphatidylglycerol phospholipid bilayers, simulated for 0.5 μs using the MARTINI force field, were also shown to favor positioning away from the bilayer center and the SWNTs made direct contact with lipid headgroups. Overall, our results suggest that SWNT position in lipid micelles is dependent on the lipid type and could potentially also be affected by the SWNT size. However, previous studies of (18,0) SWNTs encapsulated in DPPC micelles had SWNTs in micelle centers when examined in 1.6 μs long coarse-grained simulations^54^, whereas the SWNTs were asymmetrically wrapped by DPPC in brief 15 ns long atomistic simulations^62^. Therefore, it is also possible that the equilibrated structures of lipid-wrapped SWNTs may be sensitive to the type of model (atomistic or coarse-grained) used in simulations, but this possibility requires further testing and simulations.

After equilibration of POPC-SWNT conjugates, we next investigated their interactions with three known cell membrane disruptors, colistin, TAT peptide and crotamine-derived peptide. We also examined the interactions of these peptides with POPC bilayers and compared the nature and magnitude of peptide-induced perturbations in two POPC-based systems. We show that Col and TAT insert deeply into both POPC bilayers and POPC corona on SWNTs, whereas Cro only adsorbs to the POPC surface in both systems. The observed interactions agree with the previously reported mechanisms of interaction between antimicrobial cationic peptides and lipid bilayers^2,63,64^. Some antimicrobial peptides, such as for example HSP1^65^, disrupt cell membranes by adsorbing onto the lipid surface and initiating the formation of lipid-peptide particles and result in lysis of the membrane, in an antimicrobial mechanism popularly known as the carpet model^66,67^. Other antimicrobial peptides, such as Indolicidin^68^, insert into lipid bilayers and orient perpendicularly to the lipid membrane surface to generate cylindrical pores, using a mechanism known as the barrel-stave model^2^. Other peptides can generate toroidal pores^69^ or lead to spontaneous formation of pores^70^; however, pore formation processes may require exceeding threshold concentrations and cooperativity in peptide binding process^64^. In our simulations, the peptides either lie on top of the POPC bilayer/corona, which is the major binding mechanism for Cro, or insert into POPC bilayer/corona, observed for Col and TAT. Investigating other binding mechanisms would likely require longer simulations and higher peptide concentrations at POPC surfaces. Importantly, three investigated peptides bind with the same type of mechanism to both bilayers and POPC-SWNTs.

POPC-SWNT systems have one other unique interface where peptides can bind, which does not exist in POPC bilayers. This interface is the thinnest region of the POPC corona where SWNT surface is partly exposed to water. However, the peptides that induce the highest perturbation in bilayers, Col and then TAT, also induce the highest perturbations at this POPC-SWNT interface, as they lead to the increased exposure of SWNTs to water. The change in the water exposure of the SWNT surface, as well as the change in the dielectric properties of the SWNT environment, were previously shown to affect the fluorescence emission of SWNTs^43,45^.

In summary, our simulations allow us to rank the tested peptides according to the perturbation they induce in POPC-SWNT conjugates. The ranking of the peptides, from the highest to the lowest level of perturbation, is Col, TAT and Cro. The same order is observed when it comes to the number of water molecules brought by the peptides to the SWNT surface. This ranking is in agreement with the ranking of the same peptides interacting with POPC bilayers. Finally, the ranking correlates with the experimentally measured nIR optical signal change of lipid-functionalized SWNTs in response to the addition of peptides. Overall, our results indicate that lipid-functionalized SWNTs can act as simplified cell membrane models and that their optical signal transduces the peptide-induced structural perturbation of the lipid phase at the SWNT surface. These results suggest that lipid-functionalized SWNTs are promising new platforms for fast screening of new molecules that disrupt cell membranes.

## Methods

### Atomistic Models of POPC-SWNT Systems

A segment of a (6,5) single-walled carbon nanotube, 8 nm in length, was built with the *Carbon Nanostructure Builder* plugin in VMD software^71^. The structure of a single POPC molecule (1-palmitoyl-2-oleoyl-sn-glycero-3-phosphocholine) was extracted from a POPC membrane segment, built using the *Membrane Builder* plugin in VMD software and the CHARMM36 topology^72^. Then, cylindrically arranged copies of this POPC molecule were propagated around the SWNT segment using our own bash-tcl integrated script. This script arranges a user-defined number of POPC molecules radially around the SWNT, so that the molecules occupy a SWNT cross-section plane, defined by the equation *z = z*_*0*_. The radially oriented POPC molecules in the defined cross-section plane are then propagated *N* number of times along the SWNT length. The input of our script is the structure of the nanotube (pdb file), the structure of the single POPC molecule (pdb file), the angle between the neighboring radially oriented POPC molecules in the cross-section plane, and *N*, the number of times that POPC molecules in the cross-section plane are propagated along the SWNT length. Our script is available on Github with instructions for use (https://github.com/vukoviclab/POPCsensor). Using this script, four POPC-SWNT systems were prepared with the POPC:SWNT mass density ratios of 9:1, 15:1, 20:1, and 30:1 (**Figure S1**). All the POPC-SWNT systems were solvated using TIP3P water molecules. Number of molecules and sizes of the built systems are summarized in **Table S1a**.

### Atomistic Model of POPC Bilayer

A POPC bilayer, built using CHARMM-GUI membrane builder^73^ and CHARMM36 topology^72^, was prepared in a tetragonal box with 160 total lipid molecules (80 per leaflet) and 31,838 TIP3P water molecules, distributed equally over two leaflets. Numbers of lipid and water molecules were selected to keep the POPC concentration in the bilayer system similar to the POPC concentration in the 15:1 POPC-SWNT system described above. Numbers of molecules in bilayer systems are summarized in **Table S2**.

### Atomistic Models of POPC-SWNT Systems and POPC Bilayers Interacting with Membrane Disrupting Peptides

Three membrane-disrupting peptides, colistin, TAT peptide and crotamine-derived peptide, were investigated in the simulations. Col structure was prepared by completing the ChemSpider MOL structure (id=65877) with the added hydrogen atoms. Col parameters, based on the CHARMM general force field^74^ (CGenFF), were obtained using the CGenFF web interface^75,76^. TAT structure, with the sequence RKKRRQRRR, was built by extracting the residues 49 to 57 from the crystal structure of the HIV-derived TAT peptide (pdbID: 1K5K)^77^. Cro structure was built by extracting residues 1 to 14 from the crystal structure of crotamine (pdbID: 4GV5)^78^ and mutating residues 9 to 14 to obtain the final peptide with the desired sequence^16^ of YKQCHKKGGKKGSG. TAT and Cro had deprotonated negatively charged C-termini and protonated positively charged N-termini. The parameters of both peptides were based on the CHARMM36 protein force field^79^.

Next, six POPC-SWNT systems were prepared with Col, TAT and Cro molecules to examine the effects these molecules have on POPC-SWNT conjugates. Each system contained a POPC-SWNT conjugate, with either 9:1 or 15:1 POPC:SWNT mass density ratio, and ten molecules of either Col, TAT or Cro, placed randomly within 20 Å of the POPC corona (**Table S1b**). Structures of POPC-SWNT conjugates used in constructing these new systems were pre-equilibrated in 1 μs MD simulations, as described below. All the newly built systems were solvated in TIP3P water and neutralized in charge by the addition of Cl^-^ ions, which neutralized the net positive charge created by the presence of Col, TAT or Cro molecules.

Separately, three POPC bilayers were prepared with Col, TAT and Cro molecules to examine the effects these molecules have on POPC bilayers. All the bilayer systems, summarized in **Table S2**, had ten molecules of either Col, TAT, or Cro, placed randomly within 20 Å of the POPC bilayer headgroups. All the bilayer systems were solvated in TIP3P water and neutralized in charge by the addition of Cl^-^ ions.

### Molecular Dynamics Simulations

All the prepared systems were simulated using the open-source NAMD2.13^80^ package and CHARMM36 force field parameters^72,74,79^. The systems were simulated using Langevin dynamics in the isothermal-isobaric NPT ensemble, with temperature and pressure set to 310 K and 1 bar, respectively, and the Langevin coefficient, γ _Lang,_ set to 1 ps^-1^. The integration time step was set to 2 fs, and Coulomb and van der Waals non-bonded interactions were evaluated every one and two time steps, respectively, for all atoms within a 12 Å cutoff distance. The long-range Coulomb interactions were evaluated using the particle-mesh Ewald (PME) method with periodic boundary conditions^81^ applied.

Pure POPC-SWNT systems with 9:1, 15:1, 20:1, and 30:1 POPC:SWNT mass density ratios were first minimized for 20,000 steps. After minimization, solvent molecules were equilibrated for 1 ns, while the SWNTs and POPC were restrained using harmonic forces with a spring constant of 1 kcal/(mol·Å^2^). Next, the systems with 9:1, 15:1, 20:1, and 30:1 ratios were equilibrated in 1 μs, 1 μs, 0.2 μs, and 0.175 μs long production runs, respectively. In production runs, harmonic force restraints with the spring constant of 1 kcal/(mol·Å^2^) were applied to the edge atoms of SWNT segments, thus preventing these segments from thermally induced translation and rotation within the periodic unit cells and effectively creating infinitely long SWNTs extending in the direction of z-axes in the simulated systems (**Figure 1**).

Six POPC-SWNT systems with added membrane disrupting molecules **(Table S1b)** were minimized for 20,000 steps and then equilibrated for 1 ns. During minimization and equilibration, SWNT and POPC molecules were restrained using harmonic forces with a spring constant of 10 kcal/(mol·Å^2^) and 1 kcal/(mol·Å^2^), respectively. Next, the systems were equilibrated in 1 μs long production runs, with the harmonic restraints applied to the edge atoms of SWNT segments, as described above, using the spring constant of 1 kcal/(mol·Å^2^). Furthermore, to allow free movement of the disruptor molecules within the original unit cell, and to prevent their diffusion along the z-axis from the original unit cell to the neighboring periodic cells, additional restraint potential, *V*, was applied on the z-coordinate of the center of mass (COM), z_COM_, of each disruptor molecule:

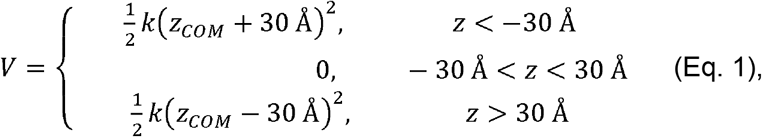

where the value of the spring constant *k* was set to 1 kcal/(mol·Å^2^).

Separately, we also examined interactions of single disruptor molecules with the equilibrated POPC-SWNT systems with 15:1 POPC:SWNT mass density ratio. In these systems, summarized in **Table S3**, one disruptor molecule of either Col, TAT, or Cro was placed above the thinnest part of the POPC corona, with the molecule COM being within ∼ 5 Å from the SWNT surface. These systems were minimized for 20,000 steps, pre-equilibrated for 1 ns, and then equilibrated in 1 μs long production runs, using the same simulation parameters and settings as defined above for the other POPC-SWNT systems. Furthermore, each production run included an additional harmonic restraint with the force constant of 10 kcal/(mol·Å^2^) applied to the COM of the disruptor molecule to keep it within 5 Å of the SWNT surface. This restraint was used to obtain long trajectories in which disruptors bind to the thinnest part of the POPC corona.

Next, we performed simulations of four POPC bilayer systems, summarized in **Table S2**. These systems include one pure POPC bilayer and three POPC bilayers with Col, TAT, and Cro molecules. The systems were minimized for 20,000 steps and then pre-equilibrated for 1 ns, while keeping disruptor molecules harmonically restrained with a spring constant of 1 kcal/(mol·Å^2^). In this pre-equilibration of bilayer systems, the pressure was maintained by keeping the ratio of the simulation box dimensions in the x-y plane constant, while allowing the fluctuations in the unit cell size along all three axes. After pre-equilibration, the systems were equilibrated in 1 μs long production runs with no restraints applied. In production runs, the pressure was adjusted by keeping the dimensions of the simulation box in the x-y plane constant and allowing fluctuations along the z-axis.

### POPC Corona Thickness Analysis

We examined the distribution of POPC around the SWNT by calculating the thickness of the POPC corona, *d*_*POPC*_, as a function of the angle *θ*, shown schematically in the inset of **Figure 1b**. The value of *d*_*POPC*_ (*θ)* for a given *θ* bin was calculated as an average over *N*_*l*_ bins along the z-axis of the system:

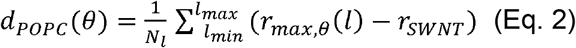

where *l*_*min*_ is the minimum z coordinate of the bin, *l*_*max*_ is the maximum z coordinate of the bin, *r*_*max,θ*_ (*l)* is the radial distance of a POPC atom which has the largest radial distance from the SWNT surface out of all the POPC atoms present in the bin, and *r*_*SWNT*_ is the radius of the (6,5) SWNT, set to the value of 0.3782 nm. Radial coordinates *r*_*max,θ*_ (*l)* and angles *θ* are defined in a cylindrical coordinate system, used because of the cylindrical shape of the SWNT, with *θ* values ranging from 0° to 360°. Values of *d*_*POPC*_ (*θ*) were calculated for every *θ* bin for the last frame of the production run trajectory (t = 1 μs), by setting the number of bins along the z-axis to *N*_*l*_ = 4 and the number of *θ* bins to 36 (with each bin spanning the angle of 10°), and defining the *l*_*min*_ and *l*_*max*_ values in the range from -22 Å to 22 Å.

To evaluate how the thinnest part of the corona evolves over time, we also calculated *d*_*POPC*_ (*θ*) over time of whole production runs and then found the minimum value of *d*_*POPC*_ (*θ*) for every time frame. The radial distances of POPC atoms for each selected time frame were calculated using a tcl script and the averaging was performed using a Python3 script.

### Calculation of Distances between Disruptor Molecules and SWNT Surfaces

To characterize the nature of binding of disruptor molecules to POPC-SWNT conjugates, we calculated the distances of all the bound disruptor molecules from SWNT surfaces. The distance of a disruptor molecule *i* from the SWNT surface averaged over time *t*, ⟨*d*_*i*_(*t*) ⟩, was calculated as:

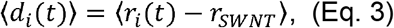

Where *r*_*i*_(*t*) is the radial distance of the center of mass of disruptor *i* at time *t*, defined in the cylindrical coordinate system, and the angle brackets denote the average over time. The time frames for which the averaging was performed were selected based on the contact criterion. Namely, if the contact area between a disruptor molecule *i* and the POPC-SWNT conjugate was greater than zero at time *t*, then the value *d*_*i*_(*t*) was considered towards the time average ⟨*d*_*i*_(*t*) ⟩. All the distances were calculated using a tcl script, and the averaging was performed using a Python3 script.

### Contact Area Calculations

To examine the extent of exposure of the SWNT surface to different molecules and functional groups, we calculated the contact areas between SWNT and different species present in the solution. Contact area between the SWNT surface and another moiety *M* at time *t, A*(*t*), was calculated as:

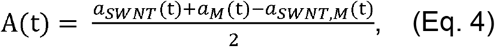

Where *a*_*SWNT*_ (*t*), *a*_*M*_(*t*) and *a*_*SWNT,M*_(*t*) are the solvent accessible surface areas (SASA) of SWNT, moiety *M*, and SWNT and moiety *M* together, at time *t*, respectively. The moieties examined include disruptor molecules, water, -NH_3_^+^ and -PO_4_^-^ zwitterionic groups of POPC, and positively charged -NH_3_^+^ groups of disruptors. The contact area calculations were performed using the SASA built-in VMD plugin where the van der Waals radius of 1.4 Å was assigned to atoms to identify the points on a sphere that are accessible to the solvent. To reduce the times required for analyses, contact areas were evaluated every 10 ns of the production run trajectories.

### Calculation of Distances between Disruptor Molecules and Centers of POPC Bilayers

To characterize the nature of binding of disruptor molecules to pure POPC bilayers, we calculated the z-axis component of distances of all the bound disruptor molecules from the center of the POPC bilayer, *Z*_*COM,bilayer*_. The z-axis component distance of a disruptor molecule *i* from the center of bilayer, averaged over time *t*, ⟨*d*_*i*_(*t*) ⟩ was calculated as:

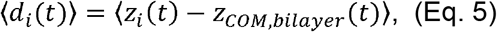

where *z*_*i*_(*t*) is the distance of the center of mass of disruptor *i* along the z-axis of the system at time *t*, and the angle brackets denote the average over time. The time frames for which the averaging was performed were selected based on how close the disruptor molecule is to the POPC bilayer. The criterion used was that if the distance between a disruptor molecule *i* and the POPC bilayer is less than 3 Å at time *t*, then the value *d*_*i*_(*t*) is counted towards the time average ⟨*d*_*i*_(*t*) ⟩. All the distances were calculated using a tcl script, and the averaging was performed using a Python3 script.

### Order Parameter Calculations

The order parameter is one of the most widely used quantities to characterize ordering in lipid membranes. It can be determined experimentally^82–84^ or calculated from atomistic MD simulations^85,86^. Here, we calculate order parameters associated with the carbon atoms forming the sn-1 and sn-2 chains of POPC molecules of the simulated bilayers. The order parameter of each carbon atom of type *i*, S_i(CH)_ is calculated as:

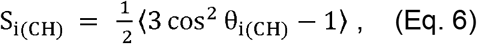

where θ_i(CH)_ is the angle between the membrane normal, which coincides with the z-axis, and a C-H bond of the carbon atom of type *i* in a selected POPC lipid chain, as shown in **Figure S7**. The angle brackets denote both ensemble and time averages, where the ensemble averaging is performed over all the carbon atoms of the same type *i* from all the POPC molecules selected for the calculation. Types of carbon atoms *i* refer to carbon atom indices in lipid chains sn-1 and sn-2, which are *i* = 1-16 and *i* = 1-18, respectively. Since POPC is an unsaturated lipid in which the sn-2 chain bears a double bond between atoms C_9_ and C_10_, the angles θ_i_ for C_9_ and C_10_ are defined as the angles which the C=C bonds make with respect to the z-axis, instead of the C-H bonds, as shown in **Figure S7**. The S_i(CH)_ values for our systems were calculated using a tcl-based script based on **Eq. 6**.

## Acknowledgments

We acknowledge the support by the NIH NIGMS-MIRA ERI grant # 1R35GM138265-01. (to N. F. R.) and the computer time provided by the Texas Advanced Computing Center (TACC).

## Notes

### Competing Interest Statement

The authors declare the following competing financial interest(s): N.E.K. and N.F. R. have an equity interest in Zymosense, Inc., a company that may potentially benefit from the research results, and also serve as company leadership. The terms of this arrangement have been reviewed and approved by Iowa State University in accordance with its conflict of interest policies.

