## Supplementary Information for "Characterizing the Interactions of Cell Membrane-Disrupting Peptides with Lipid-Functionalized Single-Walled Carbon Nanotube Systems for Antimicrobial Screening"

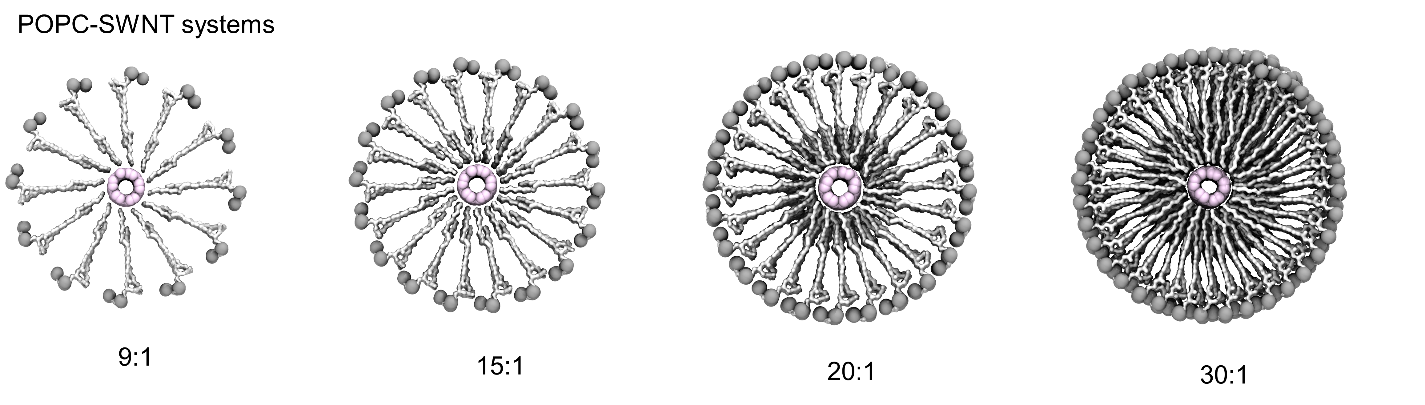

**Figure S1.** Initial structures of four POPC-SWNT systems built with POPC lipids placed in cylindrical arrangement around (6,5) SWNTs. All the systems contain a (6,5) SWNT segment of the same size (8 nm in length), and a varying number of POPC lipids, resulting in POPC-SWNT systems with POPC:SWNT mass density ratios of 9:1, 15:1, 20:1 and 30:1.

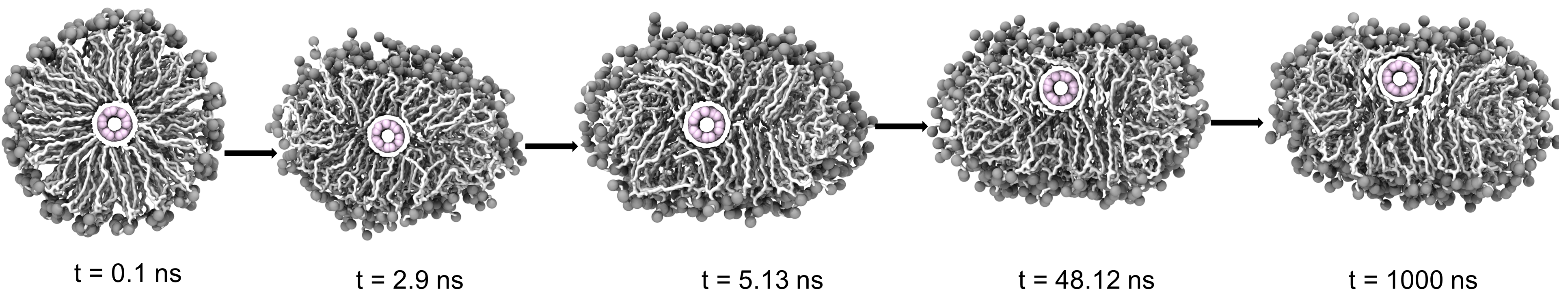

**Figure S2.** Representative snapshots of 15:1 POPC-SWNT system during a 1 μs MD simulation trajectory. Snapshots show the gradual changes in the shape of the POPC corona and the position of the SWNT in the POPC-SWNT conjugate.

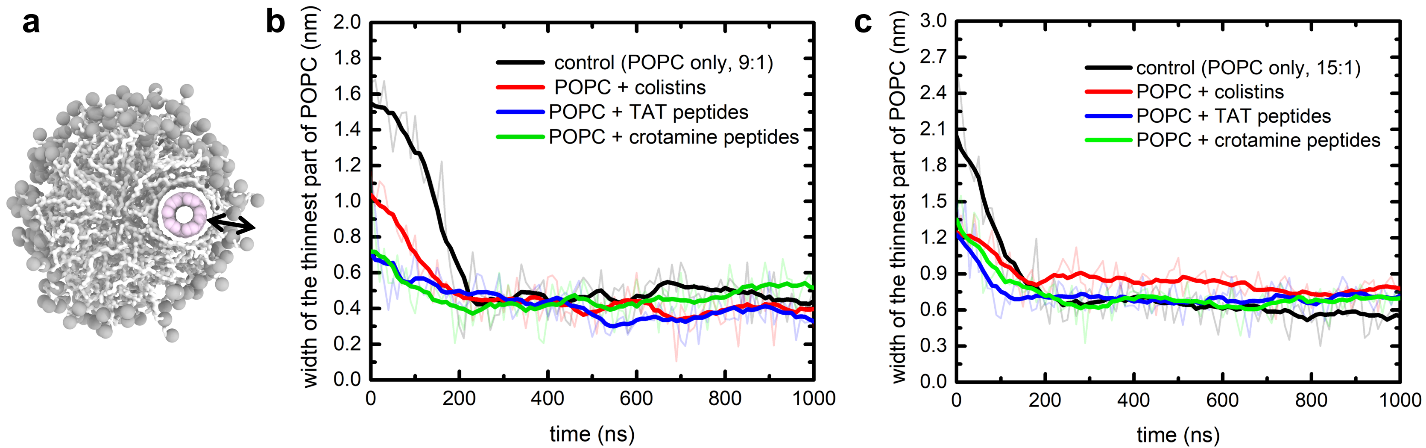

**Figure S3:** Analysis of the POPC corona asymmetry in POPC-SWNT conjugates with 9:1 and 15:1 POPC:SWNT mass density ratios. The analysis is performed by calculating the width of the thinnest part of the POPC corona over time. a) A snapshot of a representative 9:1 POPC-SWNT system, with the arrow indicating the thinnest part of the POPC corona. The color scheme is the same as in Figure 2a. Water is not shown for clarity. b) The width of the thinnest part of the POPC corona over 1 μs-long production runs for 9:1 POPC-SWNT systems. c) The width of the thinnest part of the POPC corona over 1 μs-long production runs for 15:1 POPC-SWNT systems. The lighter lines show the width of the thinnest part of the POPC corona calculated every 10 ns, and the darker lines represent the moving average. The plots in panels b) and c) are obtained by finding the minima from the analyses of the POPC corona thickness around SWNTs at individual time frames, such as the analyses shown in Figure 1b.

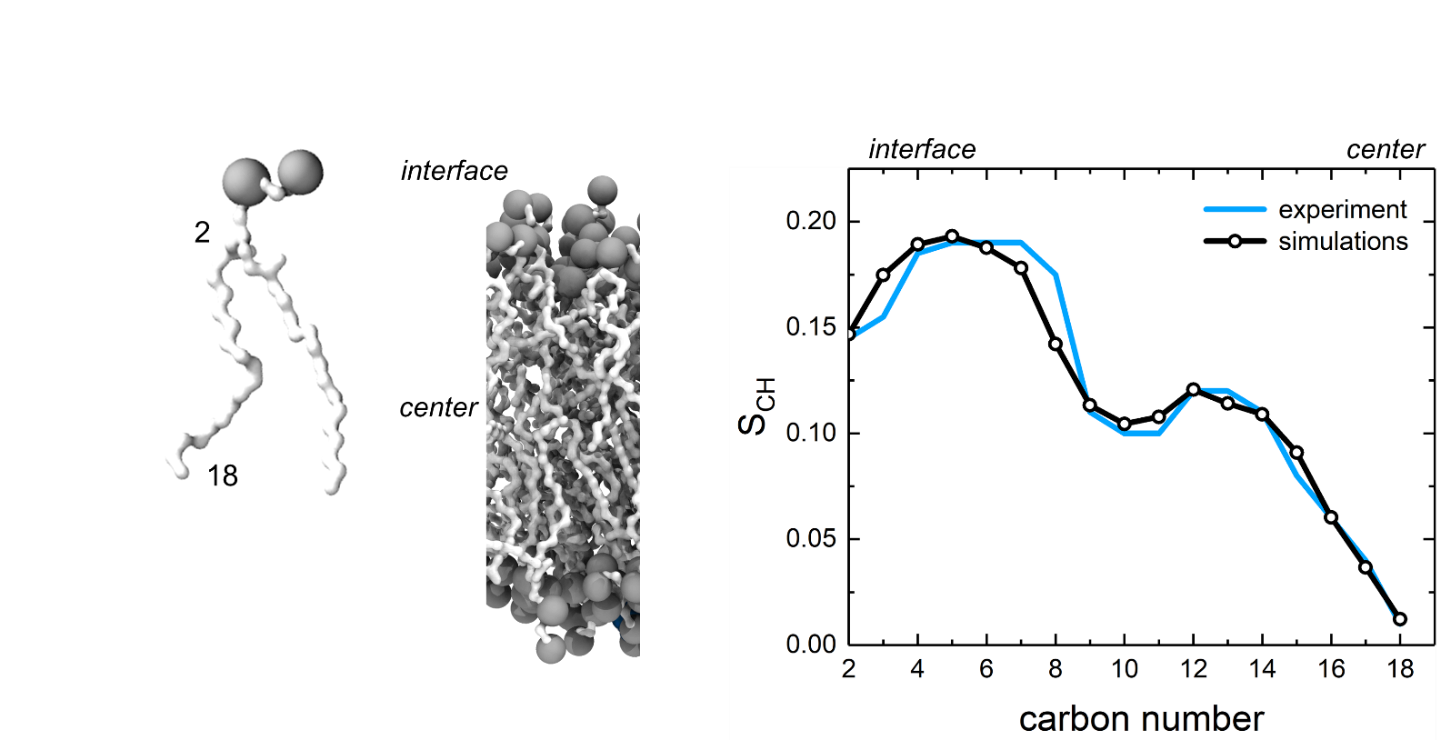

**Figure S4.** Comparison of the order parameter, S_CH_, calculated from our simulations and obtained from previous experiments^1^.

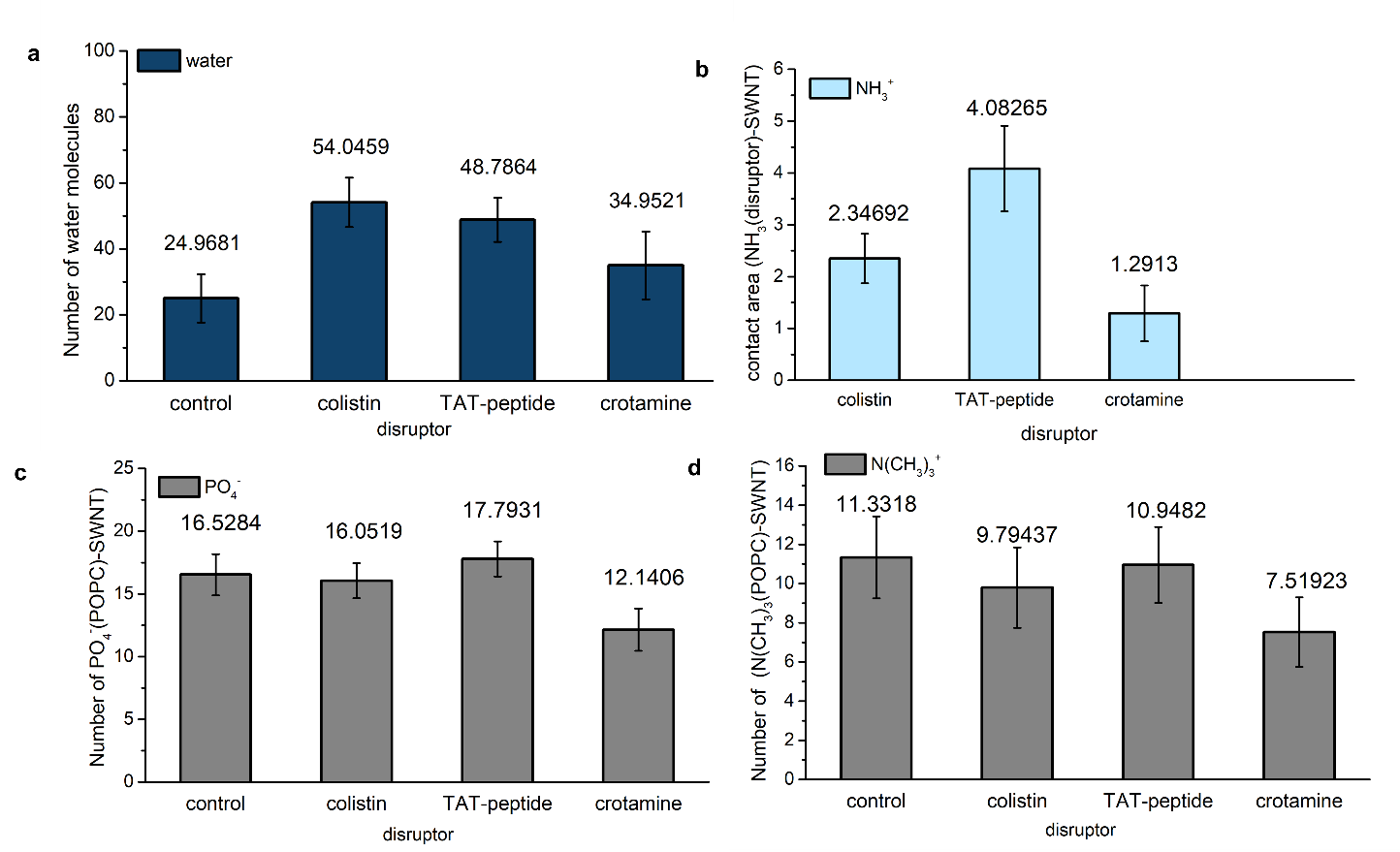

**Figure S5.** Analyses of functional groups in contact with the SWNT surface for control POPC-SWNT systems and systems interacting with Col, TAT, and Cro. a) Number of water molecules within 10 Å of the SWNT surface, which have contact area with SWNT greater than 5 Å^2^. b) Number of positively charged -NH_3_^+^ groups of disruptor molecules within 10 Å of the SWNT surface, which have contact area with SWNT greater than 5 Å^2^. c) Number of negatively charged -PO_4_^-^	groups of POPC molecules within 10 Å of the SWNT surface, which have contact area with SWNT greater than 5 Å^2^. d) Number of negatively charged -N(CH_3_)_3_^+^	groups of POPC molecules within 10 Å of the SWNT surface, which have contact area with SWNT greater than 5 Å^2^. All the numbers of molecules/groups reported in panels (a-d) are averaged over the last half of production run trajectories of the simulated systems.

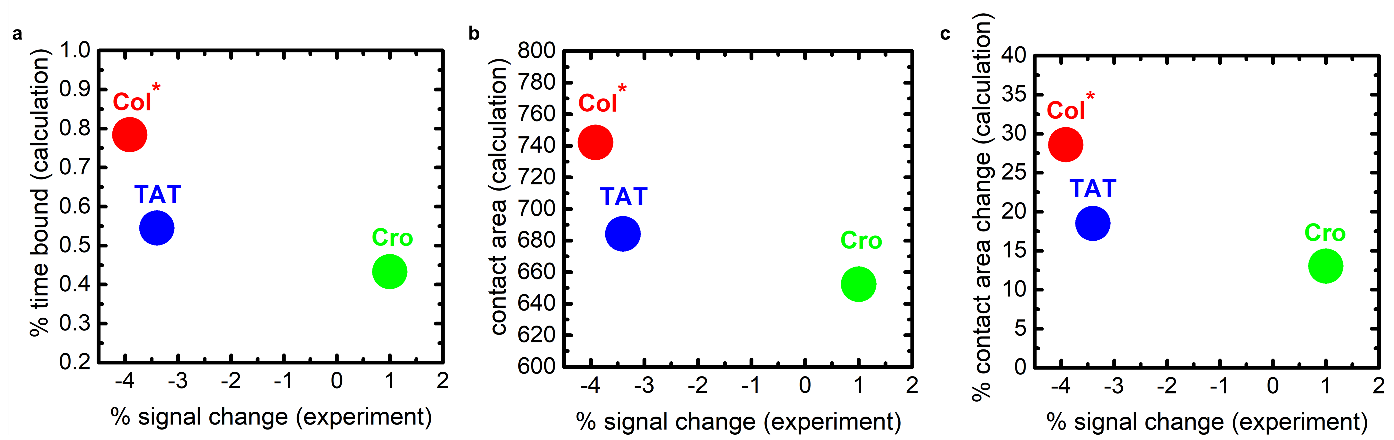

**Figure S6.** Correlation between the SWNT nIR emission signal change and three properties calculated in simulations: a) percent time that disruptors are bound to POPC-SWNT, b) the absolute water-SWNT contact area, c) percent change for water-SWNT contact area upon POPC-SWNT binding to disruptor molecules. The experimental values, reported in Ref. [2], were obtained for POPC-SWNT systems with 10 μg/mL of TAT and Cro, and for SWNT wrapped by LPS from Escherichia coli (serotype O26, Sigma L8274) with 10 μg/mL of Col.

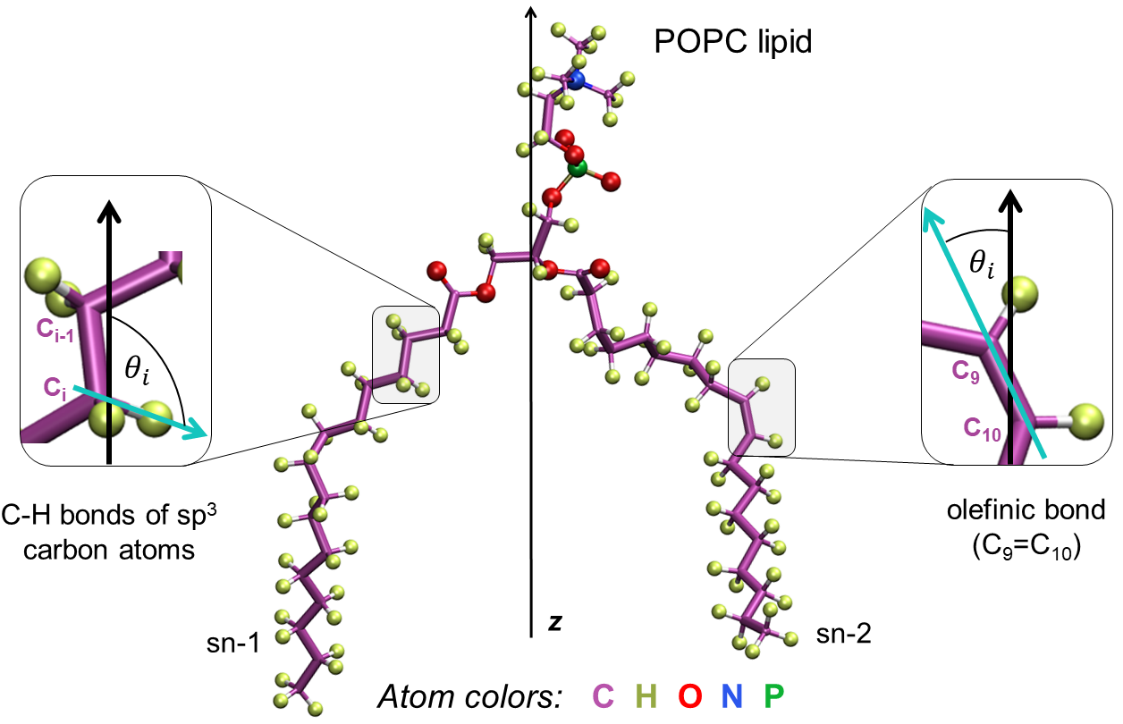

**Figure S7.** Angles used in calculation of order parameters (S_i(CH)_) for carbon atoms in POPC lipids. (left) For all sp^3^ carbon atoms on sn-1 and sn-2 chains of a POPC lipid molecule, $\theta$_i(CH)_ is the angle between the z-axis, defined as the membrane normal and shown in black color, and the line passing through the C-H bond, shown in cyan color. (right) For carbons C_9_ and C_10_ on sn-2 chain, forming the olefinic bond, $\theta$_i_ is the angle between the z-axis and the line passing through the C_9_=C_10_ bond.

**Table S1.** Summary of the simulated POPC-SWNT systems. a) Composition of POPC-SWNT control systems with varying POPC:SWNT mass density ratios. b) Composition of POPC-SWNT systems with the added membrane disruptor molecules, colistin (Col), TAT peptide (TAT), and crotamine-derived peptide (Cro).

| POPC:SWNT mass density ratio and disruptor type | | Number of POPC molecules | Number of disruptor molecules | Number of water molecules | Number of Cl^-^ ions | Size of water box ($Å^{3}$) |
| --- | --- | --- | --- | --- | --- | --- |
| **(a)**  **POPC-SWNT control systems** | | | | | | |
| 9:1 | | 96 | - | 36399 | - | 80$\times$130$\times$130 |
| 15:1 | | 161 | - | 34354 | - | 80$\times$130$\times$130 |
| 20:1 | | 215 | - | 15762 | - | 81.5$\times$93$\times$100 |
| 30:1 | | 341 | - | 15293 | - | 81.5$\times$92$\times$119 |
| **(b)**  **POPC-SWNT with ten membrane disruptors** | | | | | | |
| 9:1 | Col | 96 | 10 | 37181 | 50 | 80$\times$130$\times$130 |
|  | TAT |  | 10 | 36901 | 80 | 80$\times$130$\times$130 |
|  | Cro |  | 10 | 37026 | 50 | 80$\times$130$\times$130 |
| 15:1 | Col | 161 | 10 | 34001 | 50 | 80$\times$130$\times$130 |
|  | TAT |  | 10 | 33854 | 80 | 80$\times$130$\times$130 |
|  | Cro |  | 10 | 33760 | 50 | 80$\times$130$\times$130 |

**Table S2.** Summary of the simulated POPC bilayer systems. Composition of the systems with and without the added disruptor molecules.

| **POPC bilayer systems** | | | | |
| --- | --- | --- | --- | --- |
| System | Number of POPC molecules | Number of disruptor molecules | Number of water molecules | Number of Cl^-^ ions |
| control | 160 | - | 10613 | 0 |
| Col | 160 | 10 | 38336 | 50 |
| TAT | 160 | 10 | 38214 | 80 |
| Cro | 160 | 10 | 38094 | 50 |

**Table S3.** Summary and composition of the POPC-SWNT systems with single disruptor molecules binding to the thinnest part of the POPC corona.

| **POPC-SWNT (15:1) with single membrane disruptors** | | | | |
| --- | --- | --- | --- | --- |
| Disruptor type | Number of POPC molecules | Number of disruptor molecules | Number of water molecules | Number of Cl^-^ ions |
| Col | 161 | 1 | 35088 | 5 |
| TAT | 161 | 1 | 35028 | 8 |
| Cro | 161 | 1 | 35064 | 5 |

**Table S4.** Binding and unbinding times for disruptor molecules interacting with 9:1 and 15:1 POPC-SWNT systems. Each simulated system contained ten disruptor molecules and either 9:1 or 15:1 POPC-SWNT conjugate.

| **Col, 9:1 POPC-SWNT** | |  |  |  |  |
| --- | --- | --- | --- | --- | --- |
| **Molecule ID** | **Binding** | **Time binding starts (ns)** | **Time binding ends (ns)** | **Col-SWNT average distance and std. deviation (nm)** | **Col-SWNT contact** |
| 1 | No |  |  |  |  |
| 2 | Yes | 15 | >1000 | 2.13 (0.39) |  |
| 3 | Yes | 3 | >1000 | 3.23 (0.35) |  |
| 4 | Yes | 30 | >1000 | 2.81 (0.62) |  |
| 5 | Yes | 430 | >1000 | 1.02 (0.2) | Yes |
| 6 | Yes | 220 | >1000 | 1.36 (0.2) | Yes |
| 7 | Yes | 2 | >1000 | 1.63 (0.46) |  |
| 8 | Yes | 2 | >1000 | 2.74 (0.61) |  |
| 9 | No |  |  |  |  |
| 10 | Yes | 2 | >1000 | 1.53 (0.54) | Yes (225 ns) |
| **Col, 15:1 POPC-SWNT** | |  |  |  |  |
| **Molecule ID** | **Binding** | **Time binding starts (ns)** | **Time binding ends (ns)** | **Col-SWNT average distance and std. deviation (nm)** | **Col-SWNT contact** |
| 1 | Yes | 160 | >1000 | 2.92 (0.39) |  |
| 2 | Yes | 326.5 | >1000 | 2.89 (0.46) |  |
| 3 | Yes | 2 | >1000 | 3.23 (0.41) |  |
| 4 | Yes | 8 | >1000 | 3.35 (0.36) |  |
| 5 | Yes | 4 | >1000 | 2.06 (0.49) | Yes |
| 6 | Yes | 8 | >1000 | 3.38 (0.52) |  |
| 7 | Yes | 40 | >1000 | 3.44 (0.46) |  |
| 8 | Yes | 62 | >1000 | 3.16 (0.38) |  |
| 9 | Yes | 5 | >1000 | 2.28 (0.86) | Yes (360 ns) |
| 10 | No |  |  |  |  |

| **TAT, 9:1 POPC-SWNT** | |  |  |  |  |
| --- | --- | --- | --- | --- | --- |
| **Molecule ID** | **Binding** | **Time binding starts (ns)** | **Time binding ends (ns)** | **TAT-SWNT average distance and std. deviation (nm)** | **TAT-SWNT contact** |
| 0 | Yes | 20 | 1000 | 3.46 (0.34) |  |
| 1 | Yes | 50 | 450 | 2.26 (0.44) |  |
| 2 | Yes | 3 | 1000 | 2.28 (0.39) |  |
| 3 | Yes | 280 | 1000 | 2.63 (0.41) |  |
| 4 | No |  |  |  |  |
| 5 | Yes | 700 | 1000 | 1.31 (0.66) | Yes |
| 6 | No |  |  |  |  |
| 7 | Yes | 30 | 1000 | 1.00 (0.31) | Yes |
| 8 | Yes | 900 | 1000 | 3.27 (0.49) |  |
| 9 | Yes | 120 | 715 | 2.93 (0.45) |  |
| **TAT, 15:1 POPC-SWNT** | |  |  |  |  |
| **Molecule ID** | **Binding** | **Time binding starts (ns)** | **Time binding ends (ns)** | **TAT-SWNT average distance and std. deviation (nm)** | **TAT-SWNT contact** |
| 0 | Yes | 10, 960 | 320, 1000 | 3.49 (0.41) |  |
| 1 | Yes | 13 | 1000 | 3.94 (0.32) |  |
| 2 | No |  |  |  |  |
| 3 | Yes | 10 | 1000 | 3.42 (0.26) |  |
| 4 | Yes | 370 | 1000 | 3.40 (0.44) |  |
| 5 | Yes | 420 | 1000 | 2.98 (0.56) |  |
| 6 | Yes | 25 | 950 | 3.22 (0.39) |  |
| 7 | No |  |  |  |  |
| 8 | Yes | 400 | 1000 | 1.33 (0.28) | Yes |
| 9 | Yes | 10, 450 | 230, 1000 | 3.28 (0.48) |  |

| **Cro, 9:1 POPC-SWNT** | | |  | |  | |  | |
| --- | --- | --- | --- | --- | --- | --- | --- | --- |
| **Molecule ID** | **Binding** | **Time binding starts (ns)** | | **Time binding ends (ns)** | | **Cro-SWNT average distance and std. deviation (nm)** | | **Cro-SWNT contact** |
| 1 | Yes | 113 | | 1000 | | 2.21 (0.93) | |  |
| 2 | Yes | 220, 620 | | 415, 960 | | 3.59 (0.48) | |  |
| 3 | Yes | 180 | | 1000 | | 3.40 (0.58) | |  |
| 4 | No |  | |  | |  | |  |
| 5 | No |  | |  | |  | |  |
| 6 | Yes | 3, 470 | | 420, 600 | | 2.23 (0.86) | |  |
| 7 | Yes | 60, 800 | | 500, 1000 | | 2.93 (0.69) | |  |
| 8 | No |  | |  | |  | |  |
| 9 | Yes | 950 | | 1000 | | 1.97 (0.58) | |  |
| 10 | Yes | 673 | | 1000 | | 1.45 (1.01) | | Yes |
| **Cro, 15:1 POPC-SWNT** | | |  | |  | |  | |
| **Molecule ID** | **Binding** | **Time binding starts (ns)** | | **Time binding ends (ns)** | | **Cro-SWNT average distance and std. deviation (nm)** | | **Cro-SWNT contact** |
| 1 | Yes | 920 | | 1000 | | 3.69 (0.17) | |  |
| 2 | No |  | |  | |  | |  |
| 3 | Yes | 600 | | 1000 | | 3.97 (0.41) | |  |
| 4 | Yes | 140, 860 | | 740, 1000 | | 3.44 (0.67) | |  |
| 5 | Yes | 12, 460 | | 320, 520 | | 3.41 (0.45) | |  |
| 6 | Yes | 2 | | 1000 | | 3.47 (0.57) | |  |
| 7 | Yes | 100 | | 840 | | 3.22 (0.55) | |  |
| 8 | Yes | 5, 360 | | 185, 1000 | | 3.59 (0.49) | |  |
| 9 | No |  | |  | |  | |  |
| 10 | Yes | 40, 495 | | 240, 1000 | | 3.16 (0.95) | |  |

**Table S5.** Binding and unbinding times for disruptor molecules interacting with POPC bilayers. Each simulated system contained ten disruptor molecules and the POPC bilayer, solvated in water.

| **Col, POPC bilayer** | |  |  |  |
| --- | --- | --- | --- | --- |
| **Molecule ID** | **Binding** | **Time binding starts (ns)** | **Time binding ends (ns)** | **Col- POPC bilayer center average distance and std. deviation (nm)** |
| 1 | No |  |  |  |
| 2 | Yes | 23.1 | 1000 | 1.24 (0.59) |
| 3 | Yes | 289.9 | 1000 | 2.36 (0.81) |
| 4 | No |  |  |  |
| 5 | Yes | 8 | 61.2 | 3.42 (0.59) |
| 6 | No |  |  |  |
| 7 | Yes | 0 | 1000 | 1.88 (0.75) |
| 8 | Yes | 916 | 1000 | 0.97 (0.36) |
| 9 | No |  |  |  |
| 10 | No |  |  |  |
| **TAT, POPC bilayer** | |  |  |  |
| **Molecule ID** | **Binding** | **Time binding starts (ns)** | **Time binding ends (ns)** | **TAT- POPC bilayer center average distance and std. deviation (nm)** |
| 1 | Yes | 10 | 1000 | 2.72 (0.51) |
| 2 | Yes | 3.5,343 | 270,620 | 1.87 (0.68) |
| 3 | Yes | 55 | 1000 | 1.45 (0.80) |
| 4 | No |  |  |  |
| 5 | No |  |  |  |
| 6 | Yes | 817 | 857 | 2.22 (0.14) |
| 7 | Yes | 40 | 872.5 | 2.99 (0.62) |
| 8 | No |  |  |  |
| 9 | No |  |  |  |
| 10 | No |  |  |  |
| **Cro, POPC bilayer** | |  |  |  |
| **Molecule ID** | **Binding** | **Time binding starts (ns)** | **Time binding ends (ns)** | **Cro-POPC bilayer center average distance and std. deviation (nm)** |
| 1 | Yes | 438 | 485 | 2.39 (0.10) |
| 2 | Yes | 18,610 | 554,1000 | 2.26 (0.53) |
| 3 | Yes | 506,608 | 580.5,1000 | 1.98 (0.41) |
| 4 | No |  |  |  |
| 5 | No |  |  |  |
| 6 | Yes | 654 | 960 | 1.66 (0.36) |
| 7 | Yes | 58,139 | 125,430 | 2.35 (0.75) |
| 8 | No |  |  |  |
| 9 | No |  |  |  |
| 10 | No |  |  |  |
